# Optimization and uncertainty analysis of ODE models using 2nd order adjoint sensitivity analysis

**DOI:** 10.1101/272005

**Authors:** Paul Stapor, Fabian Fröhlich, Jan Hasenauer

## Abstract

**Motivation:** Parameter estimation methods for ordinary differential equation (ODE) models of biological processes can exploit gradients and Hessians of objective functions to achieve convergence and computational efficiency. However, the computational complexity of established methods to evaluate the Hessian scales linearly with the number of state variables and quadratically with the number of parameters. This limits their application to low-dimensional problems.

**Results:** We introduce second order adjoint sensitivity analysis for the computation of Hessians and a hybrid optimization-integration based approach for profile likelihood computation. Second order adjoint sensitivity analysis scales linearly with the number of parameters and state variables. The Hessians are effectively exploited by the proposed profile likelihood computation approach. We evaluate our approaches on published biological models with real measurement data. Our study reveals an improved computational efficiency and robustness of optimization compared to established approaches, when using Hessians computed with adjoint sensitivity analysis. The hybrid computation method was more than two-fold faster than the best competitor. Thus, the proposed methods and implemented algorithms allow for the improvement of parameter estimation for medium and large scale ODE models.

**Availability:** The algorithms for second order adjoint sensitivity analysis are implemented in the Advance MATLAB Interface CVODES and IDAS (AMICI, https://github.com/ICB-DCM/AMICI/). The algorithm for hybrid profile likelihood computation is implemented in the parameter estimation toolbox (PESTO, https://github.com/ICB-DCM/PESTO/). Both toolboxes are freely available under the BSD license.

**Supplementary information:** Supplementary data are available at *Bioinformatics* online.

## 1 Introduction

In systems and computational biology, ordinary differential equation (ODE) models are used to gain a holistic understanding of complex processes (Becker *et al.*, 2010; Swameye *et al.*, 2003). Unknown parameters of these ODE models, e.g., synthesis and degradation rates, have to be estimated from experimental data. This is achieved by optimizing an objective function, i.e. the likelihood or posterior probability of observing the given data (Raue *et al.*, 2013a). This optimization problem can be solved using multi-start local, global, or hybrid optimization methods (Raue *et al.*, 2013a; Villaverde *et al.*, 2015). Since experimental data are noise-corrupted and in most cases, only a subset of the state variables is observable, the inferred parameter estimates are subject to uncertainties. These uncertainties can be assessed using profile likelihood calculation (Raue *et al.*, 2009) and sampling (Girolami & Calderhead, 2011).

Many of the algorithms, which are applied in optimization or profile likelihood computation, exploit the gradient and the Hessian of the objective function or approximations thereof. These quantities can be used to determine search directions in optimization (Balsa-Canto *et al.*, 2001; Vassiliadis *et al.*, 1999) or to update the vector field in integration-based profile calculation (Chen & Jennrich, 1996). However, the evaluation of gradient and Hessian using standard approaches, i.e. finite differences or forward sensitivity analysis, is computationally demanding for high-dimensional ODE models. To accelerate the calculation of the objective function gradient, first order adjoint sensitivity analysis have been developed and applied (see (Fröhlich *et al.*, 2017a) and references therein). In engineering problems, similar concepts have been proposed for the calculation of the Hessian (Cacuci, 2015), but until now, these methods were never adapted to parameter estimation in biological applications.

In this manuscript, we provide a comprehensive formulation of second order adjoint sensitivity analysis for ODE constrained parameter estimation problems with discrete-time measurements. We outline the algorithmic evaluation of the Hessian and discuss the computational complexity. We include the functionality in the Advanced MATLAB Interface for CVODE and IDAS (AMICI). Furthermore, we introduce a hybrid approach for the calculation of profile likelihoods, which combines the ideas the two currently existing approaches and exploits the Hessian. We provide detailed comparisons of optimization and profile likelihood calculation of the proposed approaches and state-of-the-art methods based on published models of biological processes. Our analysis reveals that the robustness of optimization can be improved using Hessians. Moreover, we find that the hybrid method outperforms existing approaches for profile likelihood computation in terms of accuracy and computational efficiency when combined with second order adjoint sensitivity analysis.

## 2 Methods

### 2.1 Mathematical model

We consider ODE models of biological processes. The temporal evolution of a chemical concentration vector 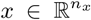 is given by a vector field *f*, depending on unknown parameters 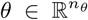 and time 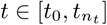:

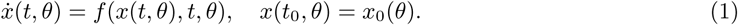

The initial state *x*_0_ may be parameter dependent. As in most applications not all states can be observed directly, a set of observables 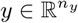 is defined:

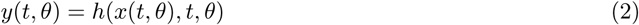

Measurements are usually noise-corrupted and this noise is modelled as normally distributed random variables with standard deviation *σ*_*ij*_ for observable *i* = 1, *…, n*_*y*_ and time point *j* = 1, *…, n*_*t*_:

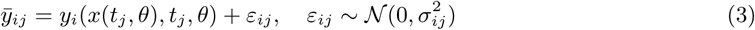

If the noise is unknown, it can be modelled as parameter dependent *σ*_*ij*_ = *σ*_*ij*_ (*θ*) and inferred from the data together with the other parameters. In the following, we will assume the *σ*_*ij*_ to be known. All derivations for parameter dependent *σ*_*ij*_ can be found in Section 1 of the Supplementary Information.

### 2.2 Parameter optimization

To infer the unknown parameters *θ*, we maximize the likelihood of observing the data given the parameter vector *θ*. Hence, the maximum likelihood estimate *θ*^*\**^ is defined as:

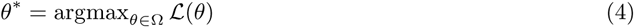

ℒ (*θ*) depends on the solution of the model and hence estimating *θ*^*\**^ is an ODE-constrained optimization problem. It must be solved numerically, since the considered ODEs rarely have closed form solutions. To improve numerical stability, we use the negative logarithm of the likelihood function as objective function, 𝒥 (*θ*) = *−ℒ* (*θ*), for minimization:

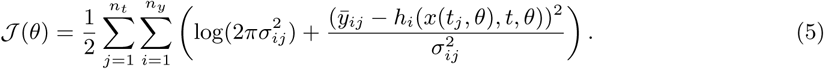

Typically, the considered optimization problems are non-convex and possess multiple local optima.

In this study, we solve the optimization problems using multi-start local optimization, an approach which has been shown to perform well in systems and computational biology (Raue *et al.*, 2013b). Initial points for local optimizations are drawn randomly from a biologically plausible region 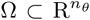 of the parameter space and the results of these optimizations are sorted by their final objective function value. Local optimization is carried out using either least-squares algorithms such as the Gauss-Newton-type methods combined with trust-region algorithms (Coleman & Li, 1996; Dennis *et al.*, 1981), or constraint optimization algorithms, which compute descent direction with (quasi-)Newton-type methods combined with interior-point or trust-region algorithms (Byrd *et al.*, 2000). Convergence of these methods can usually be improved, if the computed derivatives are accurate (Raue *et al.*, 2013b). Common leastsquares algorithms such as the MATLAB function lsqnonlin only use first order derivatives of the residuals, whereas constraint optimization algorithms like the MATLAB function fmincon exploit first and second order derivatives of the objective function.

### 2.3 Profile likelihood calculation

Since experimental data are limited, parameter estimates are subject to uncertainties. Profile likelihoods (hereafter called profiles), introduced in (Raue *et al.*, 2009), are a common method to assess these uncertainties (Kreutz *et al.*, 2013). A profile is a maximum projection of the likelihood to a chosen parameter axis: for *θ*_*k*_, *k* ∈ {1, *…, n*_*θ*_}, the profile value at *θ*_*k*_ = *c* this given by

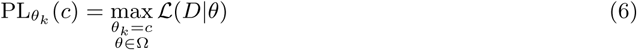

Profiles have to be computed separately for each parameter *θ*_*k*_, *k* = 1, *…, n*_*θ*_, for which currently two approaches exist.

The optimization-based approaches (as implemented in (Raue *et al.*, 2015)) computes the profile for *θ*_*k*_ via a sequence of optimization problems (Raue *et al.*, 2009). In each step, all parameters besides *θ*_*k*_ are optimized and *θ*_*k*_ is fixed to a value *c*. For each new step, *c* either increased or decreased (depending on the profile calculation direction) and the new optimization is initialized based on the previously found parameter values. As long as the function PL_*θk*_ (*c*) is smooth, this initial point will be close to the optimum and the optimization will converge within few iterations. Yet, as many optimizations have to be performed to obtain a full profile and usually all profiles have to be computed, this process is computationally demanding.

An efficient alternative to the optimization-based is the integration-based approach (Boiger *et al.*, 2016; Chen & Jennrich, 1996) (as implemented in (Kaschek *et al.*, 2016)), which circumvents the repeated optimization by using a dynamical system which evolves along the optimal path of the constraint optimization problem (6). For a constraint *g*(*θ*) = *c*, in which *g*: Ω *→* ℝ is the constraint function (in our case *g*(*θ*) = *θ*_*k*_), the dynamical system is obtained by differentiating the optimality condition

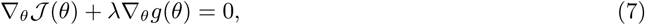

with respect to the value of the constraint, *c*, where *λ* is a Lagrangian multiplier. This yields the differential equation

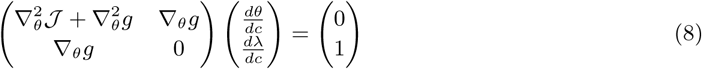

which can in principle be integrated with established differential equation solvers given the Hessian 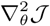 or an approximation thereof (Chen & Jennrich, 2002). However, integrating the ODE in (8) is non-trivial, as the matrix on the left hand side can degenerate and the profile path may have discontinuities. This leads to small step sizes during ODE integration. Moreover, the trajectory of the ODE solver may deviate from the true profile path of Equation (6) due to numerical errors or approximations being used.

In this study, we introduce a hybrid optimization- and integration-based approach to handle discontinuities and to ensure optimality. Our hybrid approach employs by default the integration-based approach using a high-order Adams-Bashforth scheme (Shampine & Reichelt, 1997). A pseudo-inverse is used if the matrix in (8) is degenerated. If the step size falls below a previously defined threshold, integration will be stopped and a few optimization-based steps are carried out to circumvent numerical integration problems and to accelerate the calculation. Then, integration is reinitialized at the profile path. Moreover, the remaining gradient is monitored during profile integration. If it exceeds a certain value, an optimization will be started and integration reinitialized at the profile path.

### 2.4 Computation of objective function gradient and Hessian

Providing accurate derivative information is favourable for optimization and profile computation. Yet, due to the high computational complexity, gradients are sometimes not computed and Hessians even less frequently. In this section, we recapitulate available forward and adjoint sensitivity analysis methods to, subsequently, introduce second order adjoint sensitivity analysis for the efficient computation of the Hessian for ODE models. Remark: In the following, the dependencies of *f, x, h* and their derivatives on *t, θ*, and *x* are not stated explicitly. For a detailed mathematical description of all approaches, we refer to Supplementary Information, Section 1.

#### Computation of the objective function gradient

Many state-of-the-art toolboxes compute objective function gradients using forward sensitivity analysis. When differentiating Equation (5) with respect to *θ*_*k*_, the gradient is obtained:

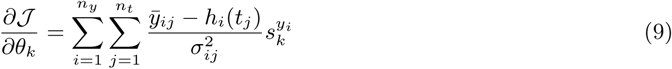

in which 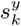 denotes the sensitivity of observable *y*_*i*_ with respect to parameter *θ*_*k*_. The observable sensitivities are calculated from the state sensitivities 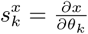 as

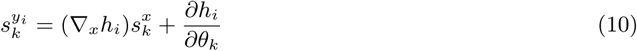

The state sensitivities need to be computed by integrating the corresponding ODE, which is obtained from differentiating Equation (1):

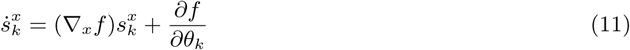

In forward sensitivities analysis, the error in the state sensitivities can be controlled together with the error of the state variables when integrating both ODEs (11) together, which makes it possible to obtain accurate gradients (Fröhlich *et al.*, 2017a). However, using this method for a system with *n*_*x*_ state variables and *n*_*θ*_ parameters requires solving an ODE of the size *n*_*x*_(*n*_*θ*_ +1). First order forward sensitivity analysis hence scales linearly in the number of parameters and in the number of state variables, which is computationally demanding for large *n*_*x*_ and *n*_*θ*_.

Adjoint sensitivity analysis circumvents the integration of the state sensitivities. In this approach, only the original ODE system (1) is integrated forward in time and subsequently the ODE for the adjoint state *p*(*t*) is integrated backward in time, starting at 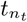:

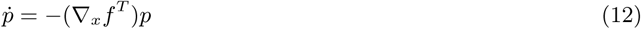

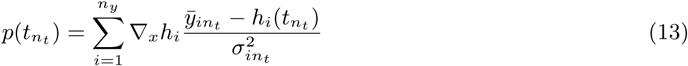

For time-discrete data, *p*(*t*) has to be reinitialized for each measurement:

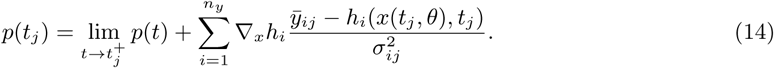

In the end, the gradient can be computed as

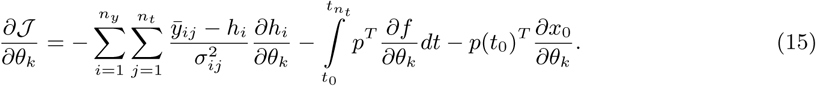

where *n*_*θ*_ one dimensional quadratures have to be computed during the backward integration. In practice, these quadratures are typically computationally less expensive, so the linear dependence of the computation time on *n*_*θ*_ for adjoint sensitivity analysis can be considered to be weak, as pointed out in (Özyurt & Barton, 2005). This yields the gradient for little more than the cost of integrating two differential equations of the size *n*_*x*_. As these scaling properties were shown to also hold true in practice (Fröhlich *et al.*, 2017a), adjoint sensitivity analysis is so far probably the most efficient method for the computation of gradients for large systems.

#### Computation and approximation of the objective function Hessian

In this study, we consider two approximations of the Hessian:

1. the Fisher Information Matrix (FIM) (Fisher, 1922)
2. the Broyden-Fletcher-Goldfarb-Shanno (BFGS) scheme (Goldfarb, 1970) and employ three approaches to compute the Hessian itself
3. central finite differences, based on gradients from adjoint sensitivities (Andrei, 2009)
4. second order forward sensitivity analysis (Vassiliadis *et al.*, 1999)
5. second order adjoint sensitivity analysis

The FIM is related to the asymptotic covariance of maximum likelihood estimates (Swameye *et al.*, 2003) and provides an approximation to the Hessian of the negative log-likelihood function. The approximation converges quadratically in the size of the residuals 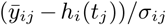 (Raue, 2013). Although, the FIM provides only an approximation, it is used in optimization, as it can be computed using first order forward sensitivities:

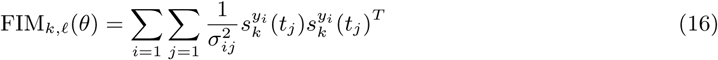

The BFGS scheme is an algorithm, which computes a positive-definite approximation to the Hessian sequentially during an optimization process using gradients, which are computed in each optimization step. Different variants of this algorithm are implemented in many state-of-the-art optimization toolboxes, like e.g. (Wächter & Biegler, 2006).

Central finite differences compute the Hessian based on perturbations in each parameter direction by a small step *δ*:

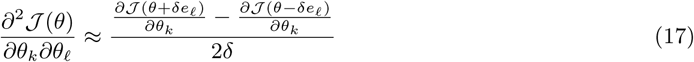

where *e*_*𝓁*_ is the unit vector with 1 at the *𝓁*-th position. The accuracy of this method depends on the step size *δ*. Good choices of *δ* depend in turn on the error tolerances of the ODE solver and are thus not easy to determine (Hanke & Scherzer (2001) and the references therein).

Second order forward sensitivity analysis extends, similar to first order forward sensitivity analysis, the considered ODE system, now including first order and second order derivatives of the state variables (see Supplementary Information, Equation (9)). If the symmetry of the Hessian is exploited, This leads to an ODE system of the size *n*_*θ*_(*n*_*θ*_ + 1)*n*_*x*_*/*2. Hence, the computational complexity of the problem scales quadratically in the number of parameters and linearly in the number of state variables, which limits this method to low-dimensional applications. Yet, second order forward sensitivity analysis yields accurate Hessians, since the error of the second order state sensitivities can be controlled during ODE integration.

Second order adjoint sensitivity analysis has so far never been applied in the field of systems and computational biology and we are not aware of any ready-to-use implementation thereof. Along the lines of first order adjoint sensitivity analysis, second order adjoint sensitivity analysis gives Hessians with better scaling properties than second order forward sensitivities. Again, the error of the Hessian can be controlled during ODE integration, yielding as accurate results as those from second order forward sensitivity analysis. To compute Hessians, the idea of the adjoint method is applied to (11) instead of (5). In a first step, the system defined by (11) is integrated forward in time. Subsequently, the corresponding adjoint system is integrated backwards in time, using the information from the forward simulation. This system consists of the original adjoint system plus the *n*_*θ*_ derivatives of *p* with respect to *θ*_*k*_.

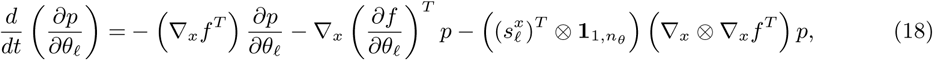

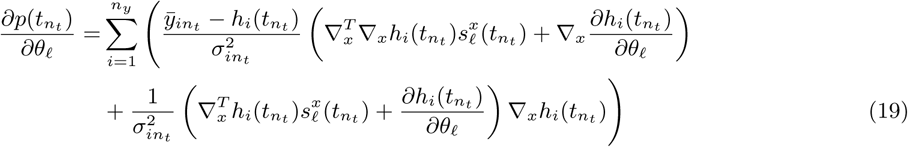

Again, the system must be reinitialized at every data time point:

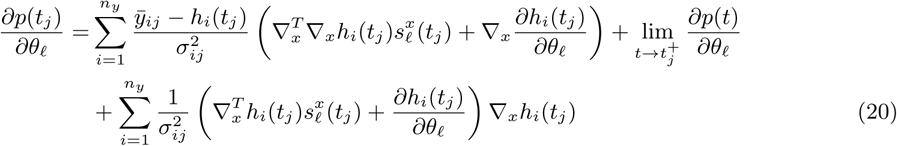

During this backward integration, 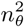 one-dimensional quadratures, which also depend on the forward trajectories of the state variables and their sensitivities, have to be calculated. Finally, the Hessian matrix can be assembled with the information coming from both ODE solves and these quadratures:

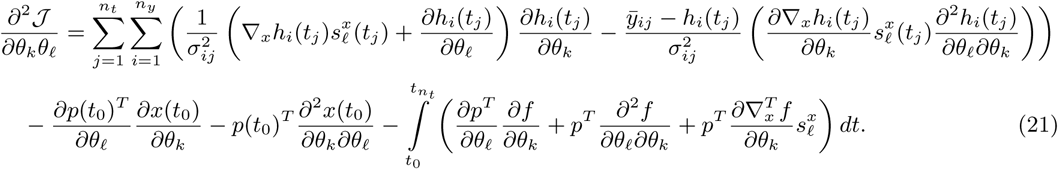

The computation of the Hessian by second order adjoint sensitivities requires solving two ODE systems of size *n*_*x*_(1 + *n*_*θ*_) and 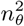 one dimensional quadratures. Again, these quadratures are fast to evaluate compared with the ODE systems. Hence, the scaling behaviour is expected to be almost linear in the number of state variables and the number of parameters.

## 3 Implementation and Results

To assess the potential of Hessian computation using second order adjoint sensitivities, we implemented the approach and we compared accuracy and computation time of the computed Hessians to to those of available methods. Furthermore, we evaluated parameter optimization and profile calculation methods using exact Hessian information for published models.

### 3.1 Implementation

The presented algorithms for the computation of gradients and Hessians by first and second order forward and adjoint sensitivity analysis were made applicable in the MATLAB and C++ based toolbox AMICI (Advanced Matlab Interface to CVODES and IDAS, Fröhlich *et al.* (2017c)), which uses the ODE solver CVODES (Serban & Hindmarsh, 2005) from the SUNDIALS package. The algorithm for hybrid profile calculation was implemented in the MATLAB toolbox PESTO (Parameter EStimation TOolbox, Stapor *et al.* (2017)).

### 3.2 Application examples

For the assessment of the methods, we considered five published models and corresponding datasets (M1 - M5). The models possess 3 to 26 state variables, 9 to 116 unknown parameters and a range of dataset sizes and identifiability properties. Four models describe signal transduction processes in mammalian cells, one describes the central carbon metabolism of E. Coli. An overview about the model properties is provided in Table 1 and a detailed description is included in the supplementary material.

**Table 1:**
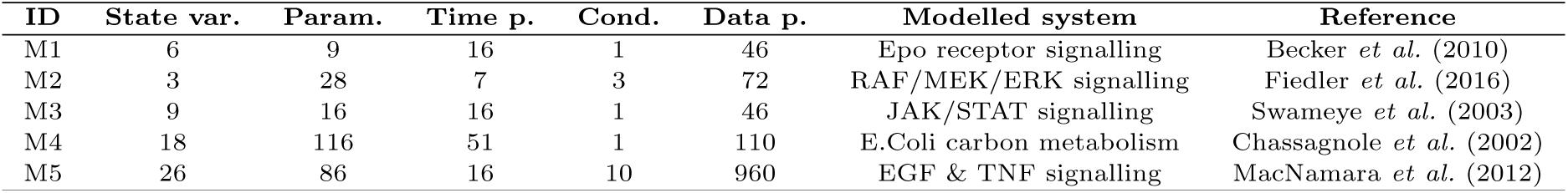
Overview of considered ODE models and their properties

| ID | State var. | Param. | Time p. | Cond. | Data p. | Modelled system | Reference |
| --- | --- | --- | --- | --- | --- | --- | --- |
| M1 | 6 | 9 | 16 | 1 | 46 | Epo receptor signalling | Becker *et al.* (2010) |
| M2 | 3 | 28 | 7 | 3 | 72 | RAF/MEK/ERK signalling | Fiedler *et al.* (2016) |
| M3 | 9 | 16 | 16 | 1 | 46 | JAK/STAT signalling | Swameye *et al.* (2003) |
| M4 | 18 | 116 | 51 | 1 | 110 | E.Coli carbon metabolism | Chassagnole *et al.* (2002) |
| M5 | 26 | 86 | 16 | 10 | 960 | EGF & TNF signalling | MacNamara *et al.* (2012) |

### 3.3 Scalability

To verify the theoretical scaling of the discussed methods, we evaluated the computation times for the model with the largest number of state variables (M5). This evaluation revealed that the practical scaling rates are close to their theoretical predictions. (Figure 1A). Second order adjoint sensitivities, Fisher information matrix and finite differences based on first order adjoint sensitivities exhibited a roughly linear scaling with respect to the parameters. Second order forward sensitivities exhibited the predicted quadratic scaling. The Fisher information matrix showed the lowest computation time for all models. The proposed approach, second order adjoint sensitivity analysis, was the fastest method to compute the exact Hessian, taking in average about 4 times as long to compute as the Fisher information matrix.

**Figure 1:**
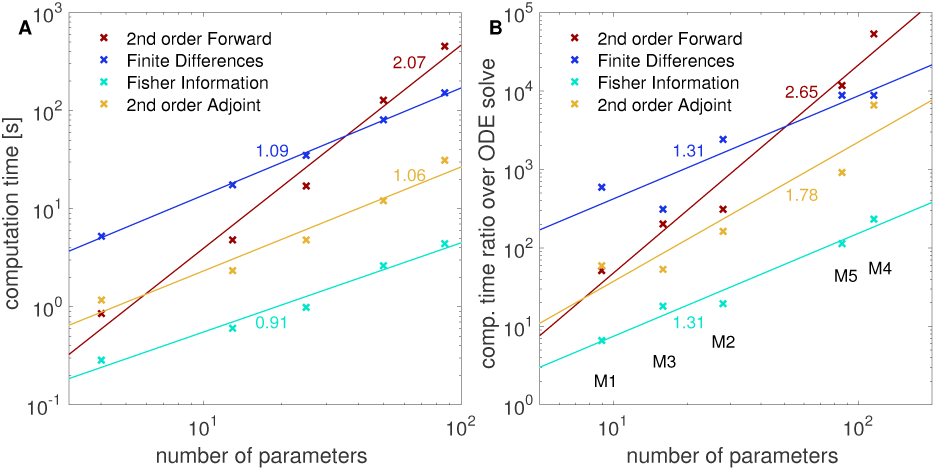
Scaling of computation times of the four investigated methods to compute or approximate the Hessian, (at global optimum for each model)including linear fits and their slopes. All reported computation times were averaged over 10 runs. A) Model (M5) was taken and the number of parameters was fixed to different values. B) The ratio of the computation times for Hessians or its approximation over the computation time for solving the original ODE is given for the five models from Table 1.

We also evaluated whether the same scaling holds across models (Figure 1B). Interestingly, we found similar but slightly higher slopes for all considered methods, although the number of state variables between models differs substantially. This suggests that in practice the number of parameters is indeed a dominating factor. Overall, second order adjoint sensitivity analysis was the most efficient method for the evaluation of the Hessian.

### 3.4 Accuracy

To assess the accuracy of Hessians and their approximations provided by the different methods, we compared the results at the global optimum. In general, we observed a good agreement of Hessians computed using second order adjoint and forward sensitivity analysis (Figure 2A). For the Hessian computed by finite difference, we found – as expected – numerical errors (Figure 2 B), which depended non-trivially on the combination of ODE solver accuracy and the step size of the finite differences. The Fisher information matrix usually differed substantially from the Hessians, even though this approximation is often considered to be good close to a local optimum (Figure 2 C).

**Figure 2:**
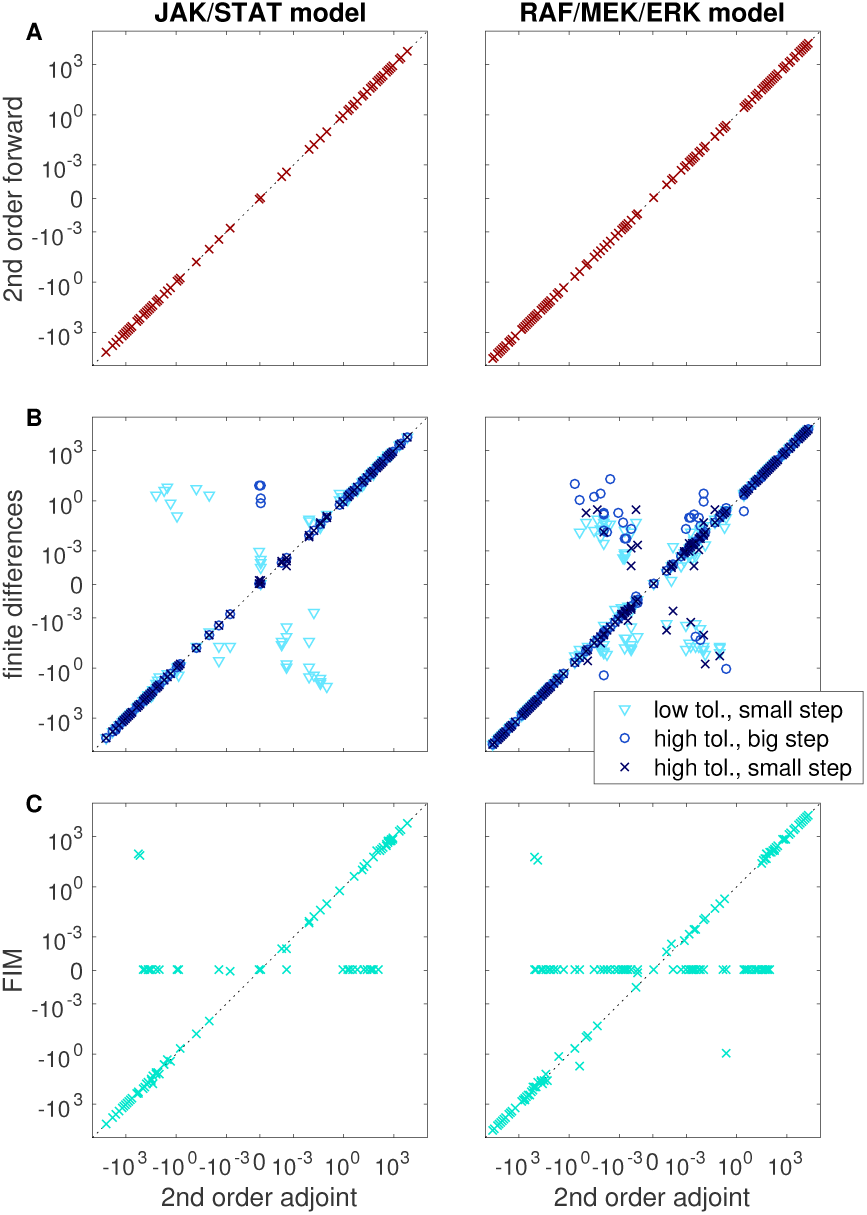
Accuracy of different methods to compute or approximate the Hessian at the global optimum for the models M2 and M3. Each point represents the numerical value of one Hessian entry as computed by two different methods: A) second order forward analysis vs. second order adjoint analysis. B) finite differences (different finite difference step sizes and ODE solver tolerances were considered) vs. second order adjoint analysis. C) Fisher information matrix vs. second order adjoint analysis. All computations were carried out with relative and absolute tolerances set to 10^−11^ and 10^−14^, respectively. For finite differences, lower accuracies of 10^−7^ and 10^−10^ were tested, together with the step sizes 10^−5^ and 10^−2^.

In combination, our assessment of scaling and accuracy revealed that second order adjoint sensitivity analysis provides the most scalable approach to obtain accurate Hessian information. Rough approximations of the Hessian in terms of the FIM could however be computed at a lower computational cost.

### 3.5 Optimization

As our results revealed an trade-off between accuracy and computation time for computation Hessians, we investigated how this affects different optimization algorithms. To this end we compared Newton and quasi-Newton variants of the interior point algorithm and the trust region algorithm:

- Residuals and their sensitivities were computed with first order forward sensitivity analysis and provided to the least-squares algorithm lsqnonlin, which used the trust-region-reflective algorithm.
- Gradient and FIM were computed using first order forward sensitivity analysis and provided to `fmincon`, which using the trust-region-reflective algorithm.
- Gradient and Hessian were computed with second order adjoint sensitivity analysis. A positive definite transformation of the Hessian was provided to `fmincon`, using the trust-region-reflective algorithm (which needs a positive definite Hessian to work).
- Gradients were calculated using first order forward sensitivity analysis and provided to `fmincon`, using the interior-point algorithm with BFGS approximation the Hessian.
- Gradient and FIM were computed with first order forward sensitivity analysis and provided to `fmincon`, using the interior-point algorithm.
- Gradients and Hessians were calculated with second order adjoint sensitivity analysis and provided to `fmincon`, using the interior-point algorithm.

The optimization study was carried out using the MATLAB toolbox PESTO for the models M2 and M3. For each of these local optimization methods, we performed four multi-start local optimizations with different initializations and 200 starting points each.

We considered the least-squares algorithm as gold standard for the considered problem class, as this method has previously been shown to be very efficient (Raue *et al.*, 2013b). Here, we studied the effect of using exact Hessians on the optimization algorithms trust-region-reflective and interior-point implemented in `fmincon`. As performance measure of the optimization methods, we considered the computation time per converged start (i.e. starts which reached the global optimum), the total number of converged starts and the number of optimization steps.

The least-square solver lsqnonlin outperformed, as expected, the constraint optimization method `fmincon` (Figure 3 and supplementary Figure 1). Among the constraint optimization methods, the methods using exact Hessians computed using the second order adjoint method, performed equal or better than the alternatives regarding overall computational efficiency (Figure 3 A). Indeed, the methods reached a higher percentage of converged starts (Figure 3 B and supplementary Figure 1) than `fmincon` using the FIM or the BFGS scheme. This is important, as convergence of the local optimizer is often the critical property (Raue *et al.*, 2013b). In addition, the number of necessary function evaluations was reduced (Figure 3 C). Furthermore, we found differences in convergence and computational efficiency for `fmincon`, depending on the chosen algorithm.

**Figure 3:**
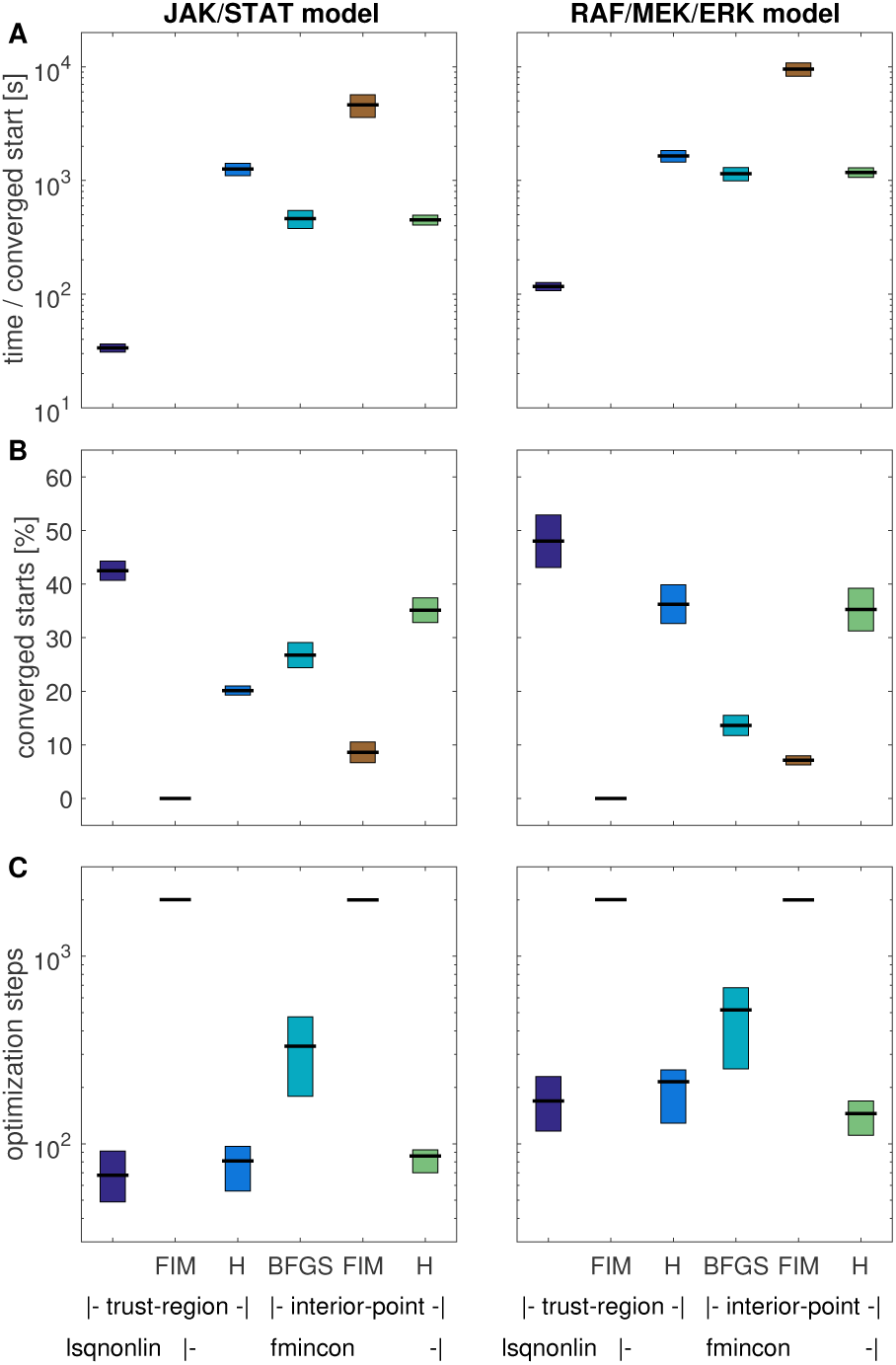
Performance measures of different local optimization methods (lsqnonlin with trust-region algorithm and `fmincon` with trust-region and interior-point algorithm, using either Hessians (H), Fisher information matrix (FIM), or the BFGS scheme). The multi-start optimization was carried out multiple times using different starting points for the local optimizations. Mean and standard deviation for A) the ratio of computation time over converged optimization starts and B) the number of converged starts are shown. C) Median and standard deviation of the number of steps over all optimization runs.

### 3.6 Profile Likelihood Calculation

To assess the benefits of Hessians in uncertainty analysis, we compared the performance of optimization- and integration-based profile calculation methods for the models M2 and M3. For the optimization based approach we employed the algorithm implemented in PESTO, which uses first order proposal with adaptive step-length selection (Boiger *et al.*, 2016). We compared the local optimization strategies described in Section 3.5 (omitting the methods based on the FIM, due to their poor performance). For the hybrid approach, we used MATLAB default tolerances for ODE integration. We compared the hybrid scheme using Hessians and the FIM. All profiles were computed to a confidence level of 95%.

The comparison of the profile likelihoods calculated using different approaches revealed substantial differences (Figure 4B and C). The optimization-based approaches worked fine for the JAK/STAT model but mostly failed for the RAF/MEK/ERK model (Figure 4A). For the RAF/MEK/ERK model, only `fmincon` with the trust-region-reflective algorithm and exact Hessians worked reliably among the optimization-based methods. Even lsqnonlin yielded inaccurate results for 11 out of 28 parameter profiles. A potential reason is that the tolerances – which were previously also used for optimization – were not sufficiently tight. Purely integration-based methods failed due to numerical problems, e.g. jumps in the profile paths. Even extensive manual tuning and the use of different established ODE solvers (including ode113, ode45, ode23, and ode15s) did not result in reasonable approximations for all profiles. In contrast, the hybrid approach provided accurate profiles for all parameters and all models, when provided with exact Hessians. When provided with the FIM, the hybrid approach failed, when it had to perform optimization, which could not rely on Hessians in this case.

**Figure 4:**
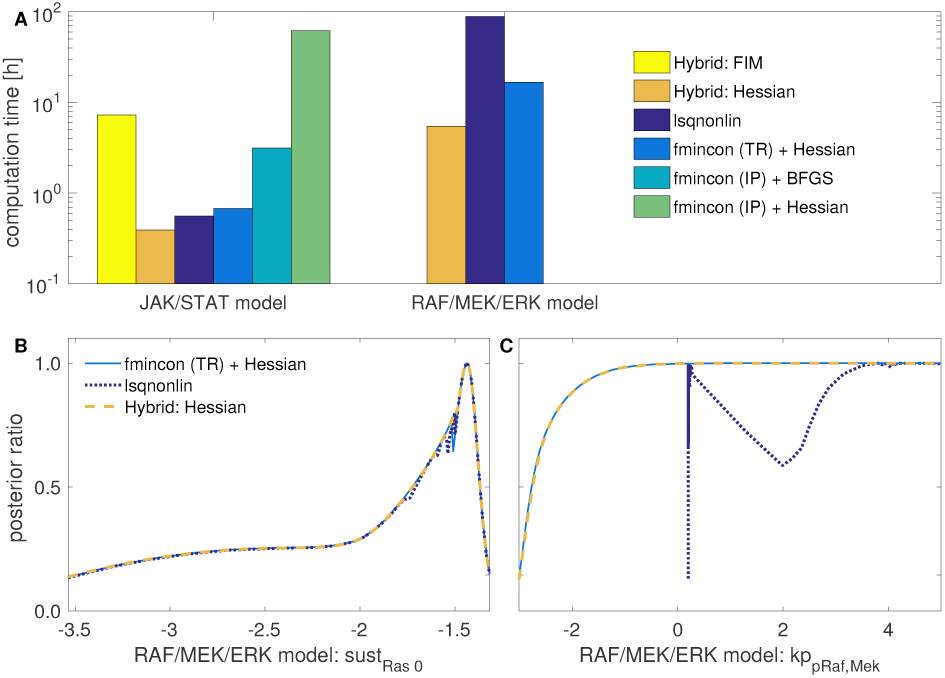
Profile likelihood computation using either the optimization-based method (`lsqnonlin` or `fmincon` with trust-region (TR) or interior-point (IP) algorithm and Hessian or BFGS approximation), or the hybrid method with either FIM or Hessian. A) Computation time for all profiles of the considered models. Three methods failed to compute profiles for the RAF/MEK/ERK model. Thus, their computation times are not depicted. B) A profile of the JAK/STAT model, all methods in good agreement with each other. C) A profile of the RAF/MEK/ERK model, lsqnonlin failed to compute the profile. D) Another profile of the RAF/MEK/ERK model, three methods in good agreement with each other.

In addition to the accuracy, also the computation time of the methods differed substantially. The hybrid method using exact Hessians was substantially faster than the remaining methods (see Figure 4A and Supplementary Information, Figure 6). The second fastest method was the optimization-based approach using the Hessian for optimization. lsqnonlin was slightly and `fmincon` using the interior-point algorithm substantially slower (for both, the BFGS scheme and Hessian), although they – as mention above – did not provide accurate profiles.

Overall, the proposed hybrid approach using exact Hessians outperformed all other methods. Compared to the best reliable competitor (optimization-based profile calculation using `fmincon` with the trust-region algorithm and exact Hessians), the computation time was reduced by more than a factor of two. This is substantial for such highly optimized routines and outlines the potential of exact Hessian information for uncertainty analysis.

## 4 Discussion

Mechanistic ODE models in systems and computational biology rely on parameter values, which are inferred from experimental data. In this manuscript, we showed that the efficiency of some of the most common methods in parameter estimation can be improved by providing exact second order derivatives. We presented second order adjoint sensitivity analysis, a method to compute accurate Hessians at low computational cost, i.e. the method scales linearly in the number of model parameters and state variables. We also provide a ready-to-use implementation thereof in the freely available toolbox AMICI.

We showed that second order adjoint sensitivity analysis possesses better scaling properties than common methods to compute Hessians while yielding accurate results, rendering it a promising alternative to existing techniques. Moreover, we demonstrated that state-of-the-art constraint optimization algorithms yield more robust results when using exact Hessians. For the computation of profile likelihoods, we demonstrated that Hessians can improve computation time and robustness of various state-of-the-art methods. Furthermore, we presented a hybrid method for profile computation, which can efficiently handle stiff and ill-conditioned problems. We also provided an implementation of this method in the parameter estimation toolbox PESTO. Although being a reliable tool in uncertainty analysis (Fröhlich *et al.*, 2014), profile likelihoods are often disregarded due to their high computational effort. The presented hybrid method based on exact Hessians is an approach the tackle this problem, as already the rudimentary implementation used in this study outperformed all established approaches.

The analysis of the optimizer performance revealed that least-squares algorithms (such as lsqnonlin), which exploit the problem structure are difficult to outperform. Many parameter estimation problems consider in systems biology do however not possess this structure. This is for instance the case for problems with additional constraints or applications considering the chemical master equation (Fröhlich *et al.*, 2016), or ODE-constrained mixture models (Hasenauer *et al.*, 2014). For these problem classes, the constraint trust-region and interior-point optimization algorithms as implemented in `fmincon` are the state-of-the-art methods. Additionally, new algorithms, which can exploit the additional curvature information, available through exact Hessian computation, in novel ways are steadily developed (see Fröhlich *et al.* (2017b)). Either directions of negative curvature can be used to escape saddle-points efficiently (Dauphin *et al.*, 2014), or third-order approximations of the objective functions are constructed iteratively from the Hessians along the trajectory of optimization to improve the convergence order (Martinez & Raydan, 2017). These approaches might outperform current optimization strategies, which are not designed to exploit directions of negative curvature that may be present in non-convex problems, and are therefore interesting subjects of further studies using the methods for Hessian computation introduced here.

While this study focused on the efficient calculation of the Hessian, second order adjoint sensitivity analysis can also be used to compute Hessian vector products. This information can be exploited by optimization methods such as truncated Newton (Nash, 1984) or accelerated conjugate gradient (Andrei, 2009) algorithms, which are suited for large-scale optimization problems. These are a few examples to illustrate how the presented results may pave the way for future improvements.

